# Resolving Inflammation: The Impact of Antiretroviral Therapy on Macrophage Traffic In and Out of the CNS

**DOI:** 10.1101/2025.05.02.651872

**Authors:** Zoey K. Wallis, Cecily C. Midkiff, Miaoyun Zhao, Soon Ok Kim, Addison Amadeck, Yiwei Wang, Maia Jakubowski, Tricia Burdo, Andrew D. Miller, Qingsheng Li, Xavier Alvarez, Robert V. Blair, Kenneth C. Williams

## Abstract

The effect of antiretroviral therapy (ART) and therapy interruption on myeloid cell traffic out of the central nervous system (CNS) with human immunodeficiency virus (HIV) and simian immunodeficiency virus (SIV) infection are understudied. Using intracisternal (i.c.) injection of dual-colored fluorescent superparamagnetic iron oxide nanoparticles (SPION) in SIV-infected macaques early (12-14 dpi) and late (30 days before sacrifice) we studied CNS macrophage viral infection, turnover, and traffic out. SPION are preferentially taken up by perivascular, meningeal, and choroid plexus macrophages. In non-infected macaques, SPION+ macrophages can traffic out of the CNS to the periphery (deep cervical lymph node (dcLN), spleen, and dorsal root ganglia (DRG)), but accumulate in the CNS with SIV infection. ART reduces the accumulation of CNS SPION+ perivascular macrophages but not meningeal or choroid plexus macrophages. ART interruption does not affect the number of SPION+ perivascular and choroid plexus but the number of SPION+ meningeal macrophages increase. ART eliminates SIV-RNA perivascular macrophages, but few scattered RNA+ macrophages in the meninges and choroid plexus remain. With ART interruption, perivascular macrophages remain SIV-but scattered SIV+ meningeal and choroid plexus macrophages exist. In non-infected animals SPION+ macrophages traffic to dcLN, spleen, and DRG at a rate that is decreased with SIV infection and AIDS. SIV- RNA+ SPION+ macrophages that traffic out of the CNS are eliminated by ART and do not rebound with ART interruption. Using two different colored SPION to study the establishment of CNS SIV viral reservoirs, we find greater numbers of early SPION+ macrophages within and outside of the CNS with SIVE, ART, and ART interruption. These data are consistent with SIV-infected perivascular macrophages establishing an early viral reservoir, and continual viral seeding of the meninges and choroid plexus during infection. These findings are consistent with ART reducing traffic of infected macrophages out of the CNS, clearing the CNS perivascular macrophage viral reservoir but not SIV-RNA+ macrophages in meninges and choroid plexus that can rebound with ART interruption.

## Introduction

Antiretroviral therapy (ART) has reduced the incidence of severe human immunodeficiency virus (HIV)-associated neurocognitive disorders (HAND) yet the prevalence of mild and asymptomatic HAND and HIV-associated neuropathy has increased (1). While durable ART diminishes HIV and simian immunodeficiency virus (SIV) infection, latently infected myeloid cells persist in the CNS and may be a source of viral recrudescence where macrophage traffic out of the CNS could result in viral redistribution (2–8). Distinct CNS macrophages, some of which originate from bone marrow (BM) monocytes, can be HIV and SIV infected and establish the CNS viral reservoir (9–11). These include perivascular macrophages, meningeal macrophages, and macrophages in the choroid plexus. CNS perivascular macrophages monitor the interface between cerebrospinal fluid (CSF) and blood (12–15), are HIV and SIV-RNA+ and DNA+ as early as 3-7 days post-infection the periphery (16, 17), and make up the CNS viral reservoir (6, 9, 10, 16). Resident meningeal macrophages, derived from mesodermal precursors in the yolk sac, are repopulated from BM at a slow, consistent rate (13, 18–21). The choroid plexus contains BM-derived stromal and Kolmer macrophages that turnover continuously(22, 23) and are HIV and SIV-RNA+ and DNA+(24–28). The resident CNS macrophage—microglia— have a minimal capacity for self-renewal and may be a viral reservoir, less so than perivascular macrophages (13, 29, 30). In normal and HIV-infected humans and SIV-infected non-human primates CNS macrophage inflammation occurs that contributes to turnover and an accumulation of BM-derived perivascular, and meningeal macrophages and CP macrophages (31–34). Less well-studied and more pressing is the effect of ART and ART interruption on these CNS macrophages with HIV and SIV infection with regards to not only their accumulation but their potential to traffic out of the CNS.

Minimally, there are three entry routes of BM and blood-derived monocytes/macrophage into the CNS including (i) migration directly from the blood via postcapillary venules in the parenchyma (perivascular macrophage), (ii) migration from postcapillary venules in the meninges (meningeal macrophage), and (iii) migration to and through the choroid plexus (stromal and Kolmer macrophage) into the CSF and subarachnoid space (SAS) (meningeal and perivascular macrophage) (35–37). Using CD34+ autologous hematopoietic stem cells transplanted in rhesus macaques, we found perivascular macrophages are continuously renewed from BM, similar to what others have shown to occur in humans and rodents(12, 14, 15, 38–42). We and others have shown early traffic (3-7 days) of virally infected monocytes/macrophages into the CNS seeds the brain with HIV and SIV (43–45) and blocking subsequent traffic with the anti-*α*4*β*1 antibody (Natalizumab) alone blocks CNS virus and resolves neuronal injury (43–45). While previous efforts have focused on blocking the migration of BM-monocytes/macrophages to the CNS, whether macrophages leave the brain and traffic out to the periphery normally and with HIV and SIV infection has not been determined. That CNS macrophage populations are reservoirs of HIV and SIV is well established but, their ability to reseed the periphery with CNS-derived virus has been suggested but not determined.

Potential pathways for exit of immune cells and fluid leaving the CNS are topics of current interest but, the dynamics of viral infection and ART, and the resolution of inflammation are understudied. Historically, this is due in part to the belief in a lack of CNS-draining lymphatics (35). Using intracisternal and intraparenchymal injection (i.c.) of dyes and fluorescently labeled antigen-presenting cells (APC), traffic of APC and drainage of CSF to the deep cervical lymph nodes (dcLN) is shown to occur normally at a rate that is increased with inflammation (19, 46–51). Ablation of meningeal lymphatics and the dcLN reduces such traffic and effectively reduces clinical symptoms in experimental autoimmune encephalomyelitis (EAE) (19, 46–51). Another potential pathway follows CSF from the fourth ventricle into the central canal and the lateral and median apertures to the SAS along the entirety of the spinal cord. This path also allows for exchange between the CSF in the central canal, CSF enveloping the spinal cord in the SAS via perivascular spaces, and is crucial for the traffic of fluid and cells out of the CNS (52, 53). While eloquent, studies using experimental injection or transplantation of labeled cells and fluid into the CNS parenchymal can result in CNS injury and/or disruption of the blood-brain barrier (BBB) and CNS parenchymal immune cell activation. Additionally, these studies have largely been done in rodents. To date, it is unknown whether macrophages leave the CNS in humans or non-human primates normally and with HIV or SIV infection. Similarly, the effects of ART on CNS macrophage viral reservoirs, CNS inflammation and macrophage traffic out, viral recrudescence, and CNS viral redistribution to the periphery are not known.

In this study we used i.c. injection of fluorescently labeled superparamagnetic iron oxide nanoparticles (SPION) to study perivascular, meningeal, and choroid plexus macrophage accumulation, traffic, and infection. We previously showed SPION preferentially labeled these macrophages in non-infected macaques in the CNS that can migrate out to secondary lymphoid organs including the dcLN, spleen, and DRG that accumulate in the CNS with SIV infection (Alverez et al., 2024 submitted). In the current study, we analyzed CNS macrophage accumulation and traffic out in SIV-infected animals with acquired immunodeficiency syndrome (AIDS) and SIV-induced encephalitis (SIVE), animals on ART, and 4 weeks post ART interruption. We find retention of SPION+ macrophages in the CNS of animals with AIDS and SIVE and decreased macrophage traffic out. ART reduces SPION+ macrophage numbers in the CNS and increases their numbers in the periphery. ART clears SIV-RNA+SPION+ perivascular macrophages but not SIV-RNA+SPION+ meningeal and choroid plexus macrophages and there is a rebound of plasma virus, SIV-RNA+SPION+ meningeal, and choroid plexus macrophages, but not CNS perivascular macrophages with ART interruption. SPION+ SIV-RNA+ macrophages are found in the periphery (dcLN, spleen, and DRG) that are cleared with ART. These findings are discussed in the context of CNS macrophage inflammation, accumulation, infection, and a viral rebound with ART interruption.

## Results

### ART reduces SPION+ perivascular macrophages but not meningeal or choroid plexus macrophages

We analyzed 3 animal cohorts; SIV-infected, that developed acquired immunodeficiency syndrome (AIDS) with SIV-induced encephalitis (SIVE) (n=5), are on ART and did not develop AIDS or SIVE (SIVnoE)(n=10) or 4 weeks after ART interruption (n=4) **(Table 1**, **Figure 1)**. Overall, there are fewer numbers of SPION+ perivascular macrophages than SPION+ meningeal macrophages (50-195x) and SPION+ choroid plexus macrophages (1-5x) **(Figure 2A-C)**. Because there are greater total numbers of macrophages in the meninges than perivascular macrophages that can take up SPION, we determined the ratio of total CD163+ and CD68+ SPION+ macrophages to total CD163+ and CD68+ macrophages in the perivascular, meningeal, and choroid plexus macrophages compartments (Perivascular SIVE 1:0.05; ART 1:0.04; ART Off 1:0.02)(Meningeal SIVE 1:0.31; ART 1:0.41; ART Off 1:0.56)(Choroid Plexus SIVE 1:0.02; ART 1:0.11; ART Off 1:0.02)**(Figure 2)**. Importantly, the ratio of SPION+ to total macrophages did not change within compartments between treatment groups. Animals with AIDS and SIVE have 0.66 ± 0.13 SPION+/mm^2^ SPION+ perivascular macrophages that were reduced (p=0.054) to 0.18 ± 0.02 SPION+/mm^2^ with ART and did not change with ART interruption (0.21 ± 0.20 SPION+/mm^2^) **(Figure 2A)**. SIVE animals had 28 ± 5.8 SPION+/mm^2^ SPION+ meningeal macrophages and no decrease with ART (22 ± 4.9 SPION+/mm^2^) and significantly (p<0.05) increased numbers with ART interruption compared to ART (ART Off: 39 ± 8.6 SPION+/mm^2^) **(Figure 2B).** There were 18 ± 13 SPION+/mm^2^ SPION+ macrophages in the choroid plexus of SIVE animals that slightly but not significantly increased with ART (26 ± 25 SPION+/mm^2^) and significantly decreased with ART interruption (8.0 ± 3.7 SPION+/mm^2^) (p<0.05) **(Figure 2C).** This decrease in the number of SPION+ macrophages in the choroid plexus following ART interruption may correspond with the increased numbers of SPION+ macrophages in the meninges as a result of macrophage traffic into the CNS. Overall, these data are consistent with the accumulation of perivascular macrophages with SIVE that is reduced with ART and an accumulation of SPION+ meningeal and choroid plexus macrophages that are unaffected by ART.

**Figure 1.**
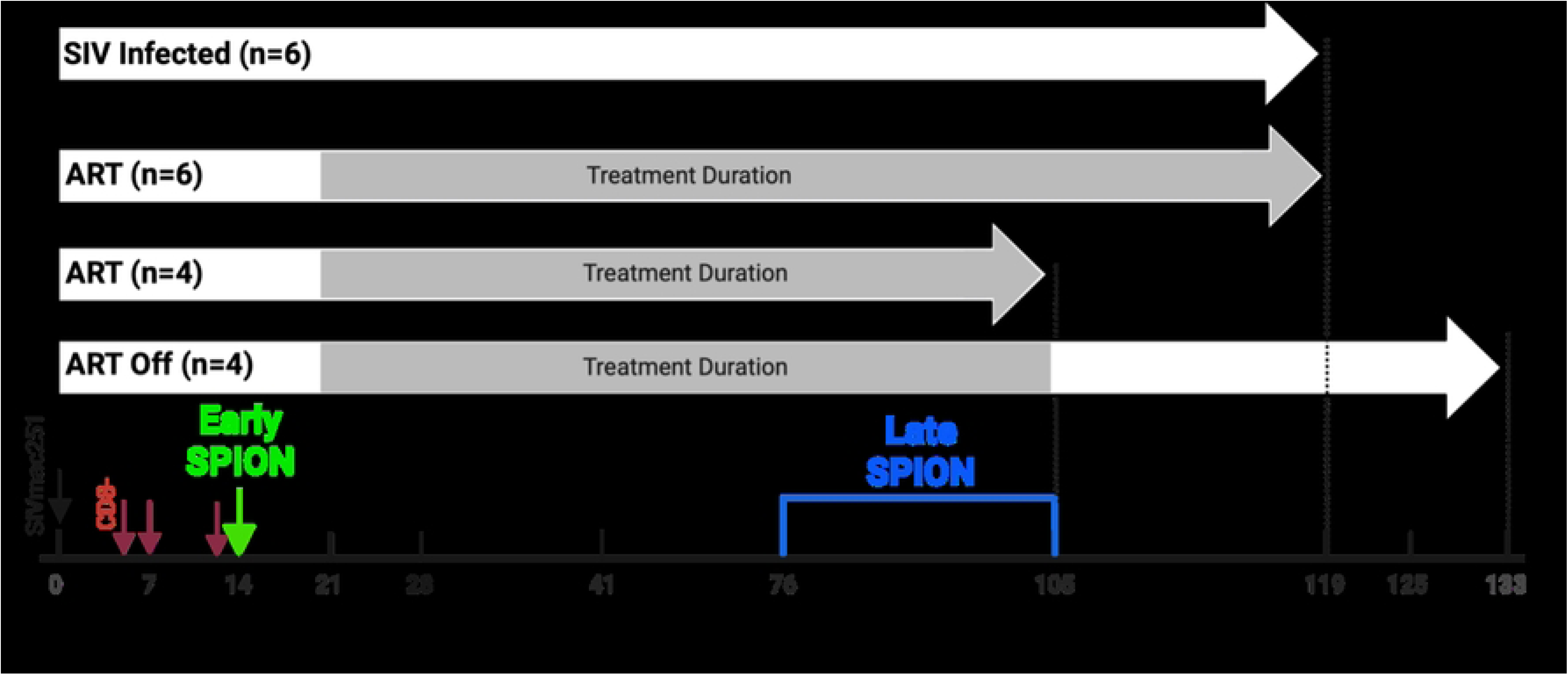
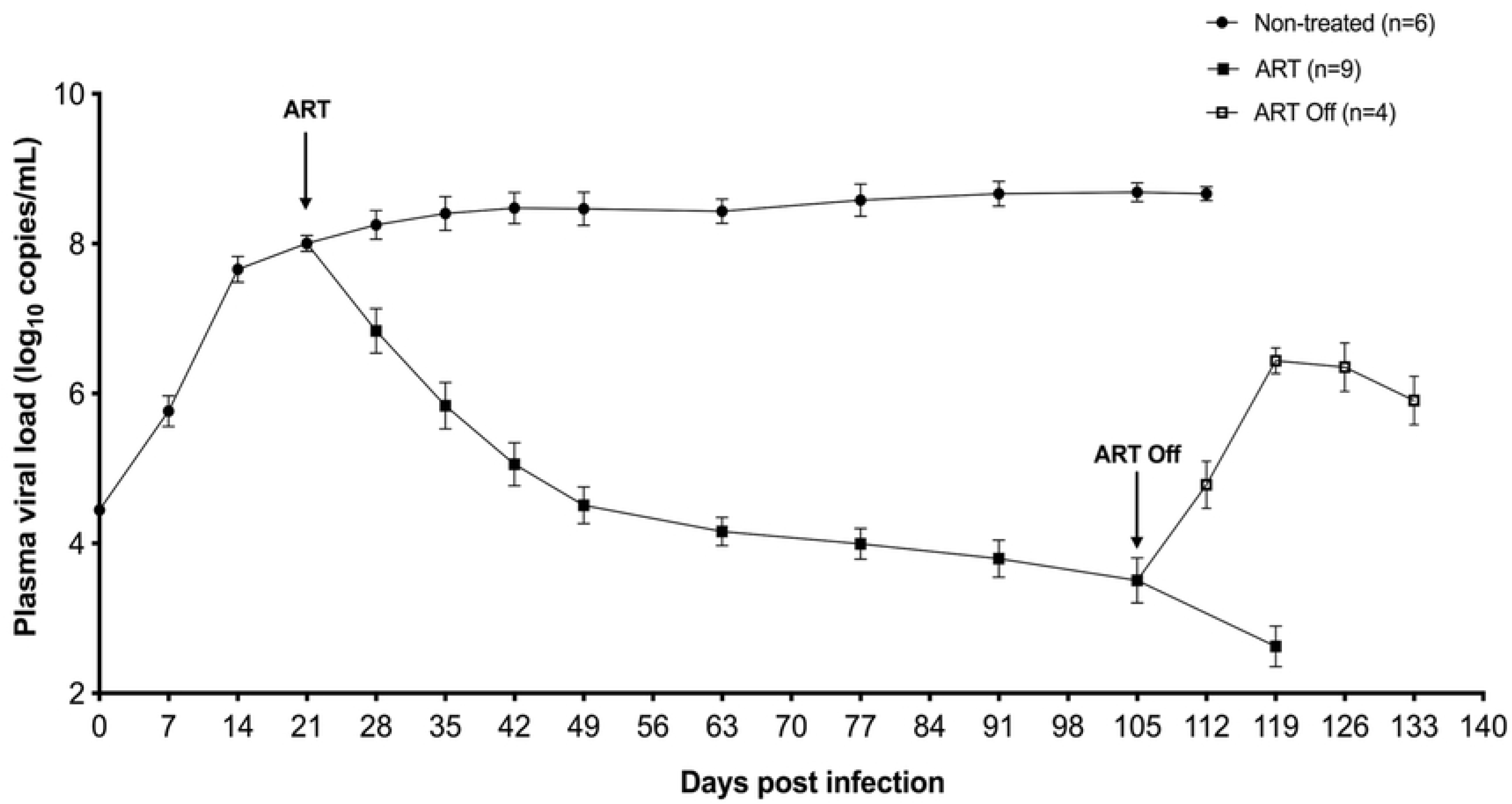
Study design and plasma viral load. **(A)** Nineteen rhesus macaques were infected with SIVmac251 (black arrow) and CD8 T lymphocytes depleted on 5, 7, and 12 dpi (red arrows). Fourteen macaques were treated with ART twice daily starting 21 dpi until 105 – 119 dpi where animals were sacrificed on ART treatment (n=10) or had ART interruption (n=4) for 4 weeks before sacrifice. Macaques received i.c. SPION injection early (12-14 dpi, Green Early SPION) and late (30 days prior to sacrifice, Late SPION). All untreated macaques were sacrificed upon progression to AIDS between 83–126 dpi. Study design created using BioRender. **(B)** Plasma viral load was measured longitudinally in SIV-infected animals (n=6), SIV-infected animals with ART (n=9), and SIV-infected with ART and following ART Off (n=4). Prism was used for graphing and statistical analysis using Kruskal-Wallis nonparametric test followed by Dunn’s post-hoc test, with significance accepted at p <0.05.

**Figure 2.**
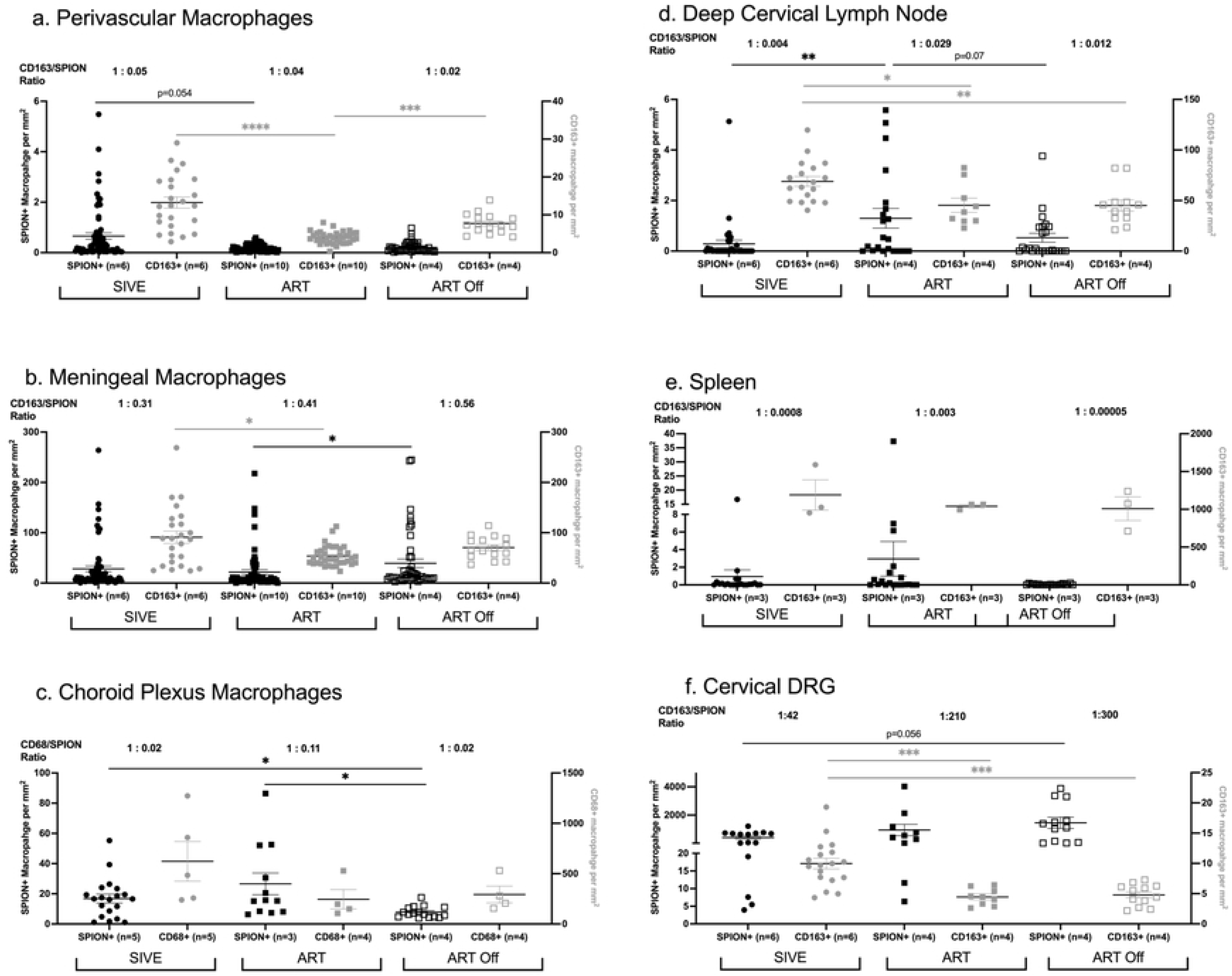
ART reduces the number of SPION+ perivascular macrophages and increases traffic out. Single-label immunohistochemistry for CD163 with Prussian Blue staining for SPION (black) or macrophage alone (grey) in non-treated SIV-infected animals with encephalitis (SIVE), SIV-infected macaques sacrificed on ART (ART), and SIV-infected macaques with 4 weeks of ART interruption (ART Off) in **(A)** perivascular space, **(B)** meninges, **(C)** choroid plexus, **(D)** dcLN, **(E)** spleen, and **(F)** DRG. Each data point represents the cell count from the cortical tissue section examined and is expressed as the number of positive cells per mm^2^. The numbers above each graph are the average ratio of CD163+SPION+/ CD163+ macrophages or CD68+SPION+/CD68+ in the choroid plexus. Prism was used for graphing and statistical analysis Kruskal-Wallis nonparametric test was followed by Dunn’s post-hoc test, ∗P < 0.05, ∗∗P < 0.01, and ∗∗∗∗P < 0.0001.

**Table 1.**
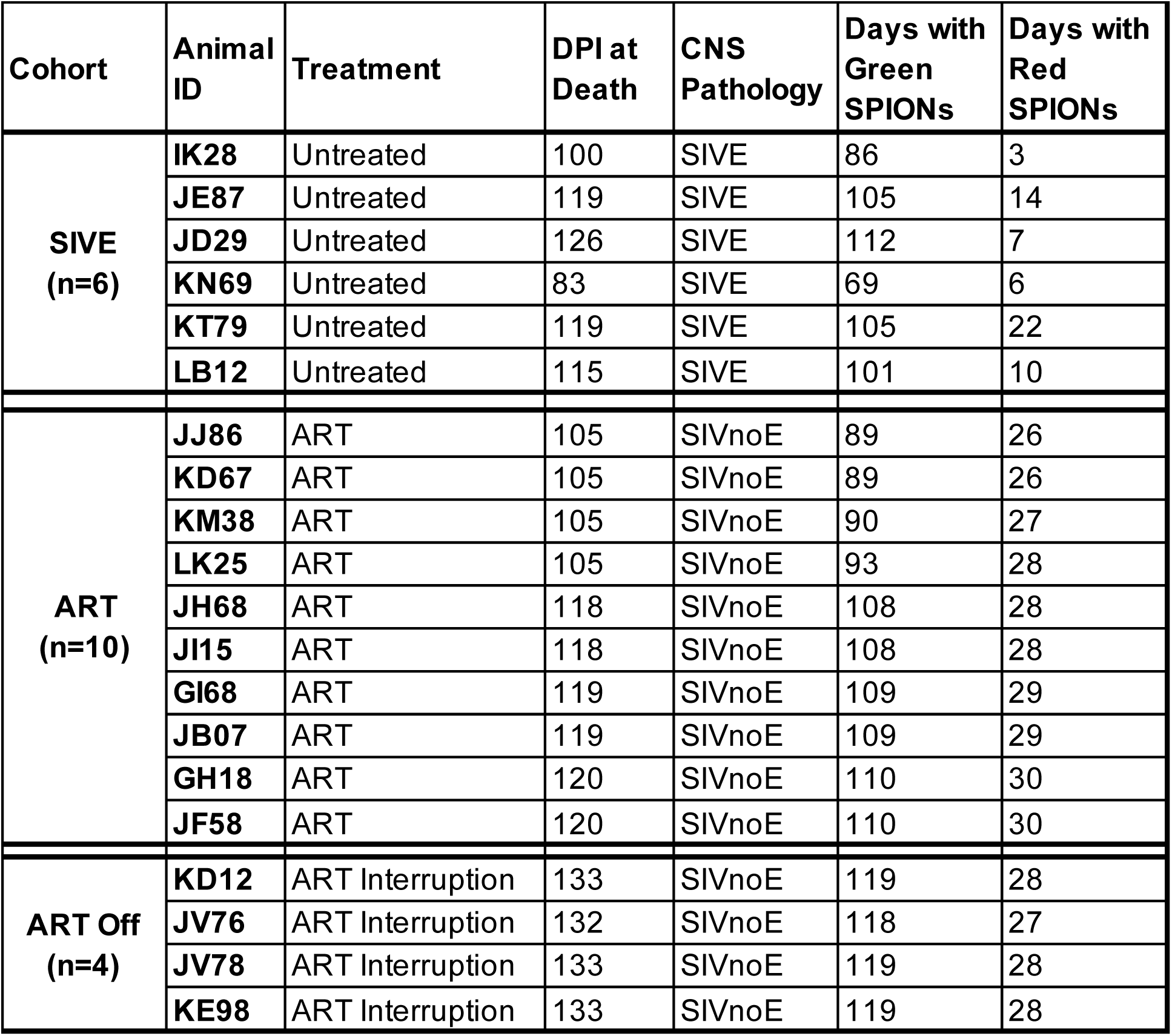
Animals Used in Study.

| Cohort | Animal ID | Treatment | DPI at Death | CNS Pathology | Days with Green SPIONs | Days with Red SPIONs |
| --- | --- | --- | --- | --- | --- | --- |
| **SIVE**<br>**(n=6)** | **IK28** | Untreated | 100 | SIVE | 86 | 3 |
|  | **JE87** | Untreated | 119 | SIVE | 105 | 14 |
|  | **JD29** | Untreated | 126 | SIVE | 112 | 7 |
|  | **KN69** | Untreated | 83 | SIVE | 69 | 6 |
|  | **KT79** | Untreated | 119 | SIVE | 105 | 22 |
|  | **LB12** | Untreated | 115 | SIVE | 101 | 10 |
| **ART**<br>**(n=10)** | **JJ86** | ART | 105 | SIVnoE | 89 | 26 |
|  | **KD67** | ART | 105 | SIVnoE | 89 | 26 |
|  | **KM38** | ART | 105 | SIVnoE | 90 | 27 |
|  | **LK25** | ART | 105 | SIVnoE | 93 | 28 |
|  | **JH68** | ART | 118 | SIVnoE | 108 | 28 |
|  | **JI15** | ART | 118 | SIVnoE | 108 | 28 |
|  | **GI68** | ART | 119 | SIVnoE | 109 | 29 |
|  | **JB07** | ART | 119 | SIVnoE | 109 | 29 |
|  | **GH18** | ART | 120 | SIVnoE | 110 | 30 |
|  | **JF58** | ART | 120 | SIVnoE | 110 | 30 |
| **ART Off**<br>**(n=4)** | **KD12** | ART Interruption | 133 | SIVnoE | 119 | 28 |
|  | **JV76** | ART Interruption | 132 | SIVnoE | 118 | 27 |
|  | **JV78** | ART Interruption | 133 | SIVnoE | 119 | 28 |
|  | **KE98** | ART Interruption | 133 | SIVnoE | 119 | 28 |

Twenty SIV-infected, CD8+ T-lymphocyte-depleted rhesus macaques were used in this study. Sections of cortical CNS tissue were examined blindly by a veterinary pathologist and the presence of SIV-induced encephalitis (SIVE) was defined by productive viral replication, the presence of MNGC, and macrophage accumulation or the presence of SIV with no encephalitis (SIVnoE). All macaques received i.c. SPION injection early (12-14 dpi, Green Early SPION) and late (26-30 days before sacrifice, Red Late SPION). ART animals began treatment daily at 21 dpi and four animals had ART interrupted (ART Off) for 4 weeks prior to sacrifice.

### ART decreases total perivascular, meningeal, and choroid plexus macrophage numbers

We next assessed the effects of ART and ART interruption on the accumulation of total macrophages in the perivascular space, meninges, and choroid plexus compartments. SIVE animals have 13 ± 1.5 CD163+/mm^2^ CD163+ perivascular macrophages that significantly decrease with ART (4 ± 0.26 CD163+/mm^2^,p<0.0001) and significantly increase with ART interruption (7.7 ± 0.66 CD163+/mm^2,^ p<0.001) **(Figure 2A)**. Similarly, ART significantly (p<0.0001) decreases the number of meningeal and choroid plexus macrophages (Meninges SIVE: 91 ± 12 CD163+/mm^2^; ART: 54 ± 3.3 CD163+/mm^2^) (Choroid plexus SIVE: 622 ± 441

CD68+/mm^2^; ART: 246 ± 192 CD68+/mm^2^) **(Figure 2B-C)**. Following ART interruption, there is a 1.5-fold increase in the number of meningeal macrophages compared to ART that did not reach statistical significance and no increase in the number of choroid plexus macrophages (meninges ART: 54 ± 3.3 CD163+/mm^2^; ART Off: 70 ± 5.5 CD163+/mm^2^) (choroid plexus ART: 246 ± 192 CD68+/mm^2^; ART Off: 293 ± 167 CD68+/mm^2^) **(Figure 2B-C)**. This suggests there is an increased accumulation of perivascular, meningeal, and choroid plexus macrophages with SIV infection and AIDS that is reduced with ART and does not significantly rebound in the meninges and choroid plexus but increases in perivascular macrophages following 4 weeks of ART interruption.

### There is increased traffic of SPION+ macrophages out of the CNS with ART

Following i.c. SPION inoculation, we found SPION+ macrophage in the optic nerve, nasal septum, cribriform plate, and in the dcLN, spleen, and dorsal root ganglia (DRG) (Wallis et al., 2025 *accepted AJP*). In contrast to the CNS and consistent with our previous findings, there are fewer SPION+ macrophages in the periphery of animals with AIDS and SIVE than animals on ART (dcLN SIVE: 0.28 ± 0.15 SPION+/mm^2^; ART: 1.3 ± 0.39 SPION+/mm^2^) (spleen SIVE: 0.95 ± 3.5 SPION+/mm^2^; ART: 3.0 ± 8.6 SPION+/mm^2^) (DRG SIVE: 420 ± 104 SPION+/mm^2^; ART: 947 ± 400 SPION+/mm^2^) **(Figure 2D-F)**. There is a trend of decreased numbers of SPION+ macrophages in the periphery following ART interruption (dcLN ART Off: 0.53 ± 0.18 SPION+/mm^2^) (spleen ART Off: 0.051 ± 0.070 SPION+/mm^2^) **(Figure 2D&E)**. The number of SPION+ macrophages in the DRG trend to increase with ART (2.3x) and with ART interruption (3.5x) compared to animals with AIDS and SIVE although they did not reach statistical significance (DRG ART: 947 ± 400 SPION+/mm^2^; ART Off: 1458 ± 398 SPION+/mm^2^) **(Figure 2F)**. Although the number of SPION+ macrophages did not significantly increase (p=0.06) in the DRG with ART interruption, there was a reduced ratio of CD163+ to CD163+SPION+ macrophages in the DRG (SIVE 1:42, ART 1:213, ART Off 1:304), suggestive of an increased in SPION+ macrophage leaving the CNS via the DRG with ART interruption. These data are consistent with SPION+ macrophages that are retained in the CNS with SIV-infection and inflammation, and increased CNS macrophages trafficking out to the periphery with ART, and resolution of CNS inflammation.

### ART decreases the number of total macrophages in the periphery

In parallel with studies of SPION+ macrophages in the periphery, we assessed the number of macrophages in the same tissue with ART and ART interruption. In the dcLN, SIVE animals have 69 ± 4.8 CD163+/mm^2^ CD163+ macrophages that significantly decrease with ART (45 ± 7.0 CD163+/mm^2^, p<0.05) and that are similar to ART after ART interruption (ART Off 45 ± 5.6 CD163+/mm^2^) **(Figure 2D)**. Similarly, ART reduces macrophages in the spleen and significantly (p<0.005) decreases macrophages in the DRG compared to animals with AIDS and SIVE (Spleen SIVE: 1190 ± 348 CD163+/mm^2^; ART: 1041 ± 40 CD163+/mm^2^) (DRG SIVE: 10 ± 0.89 CD163+/mm^2^; ART: 4.5 ± 0.45 CD163+/mm^2^) **(Figure 2E&F)**. There are no differences in the number of macrophages in the spleen and DRG between the ART and ART interruption cohorts (Spleen ART: 1041 ± 40 CD163+/mm^2^; ART Off 1008 ± 270 CD163+/mm^2^) (DRG ART: 4.5 ± 0.45 CD163+/mm^2^; ART Off 4.9 ± 0.51 CD163+/mm^2^) **(Figure 2E&F)**. These data demonstrate an increased accumulation of macrophages in the dcLN, spleen, and DRG with SIV infection and AIDS that is reduced with ART and does not rebound with ART interruption.

### ART reduces SPION+ and total SIV-RNA+ macrophages and gp41+ cells in the CNS and periphery

Using SIV-RNAscope and immunohistochemistry for SIV-RNA and SIV-gp41+ cells we counted the number of SIV-productively infected cells with SPION in the CNS and assessed the effects of ART and ART interruption **(Table 2**. **Figure 3)**. Overall, AIDS and SIVE animals have large numbers of SIV-RNA+ and gp41+ macrophages with and without SPION in the CNS and periphery that are significantly reduced with ART **(Table 2**. **Figure 3)**. Within the CNS, there are greater numbers of SIV-RNA+SPION+ meningeal (25x) and choroid plexus macrophages (3.2x) versus perivascular macrophages in AIDS and SIVE animals **(Table 2**. **Figure 3)**. There are significantly increased numbers of vRNA+ and gp41+ macrophages, including MNGC, localized primarily in CNS lesions—some of which contain SPION—and SIV-DNA+ cells (data not shown) also primarily in lesions **(Table 2**. **Figure 4).** With ART, there is a significant reduction of SIV- RNA+ and gp41+ perivascular macrophages (CD163+ p<0.005; CD68+ p<0.0001), macrophages in the meninges (CD163+ p<0.005; CD68+ p<0.005), and choroid plexus (CD68+ p=0.17) **(Table 2)**. ART fully eliminates SIV-RNA+SPION+ perivascular macrophages while few scattered SIV- RNA+SPION+ meningeal (1 SPION+SIV-RNA+ macrophage) and choroid plexus macrophages (10 SPION+SIV-RNA+ macrophages) persist **(Figure 3**. **Table 2)**. With ART interruption, there are few scattered SPION+SIV-RNA+ macrophages and gp41+ cells in the meninges, but not SPION+SIV-RNA+ or SIV-RNA+ perivascular macrophages or in the choroid plexus **(Table 2).**

**Figure 3.**
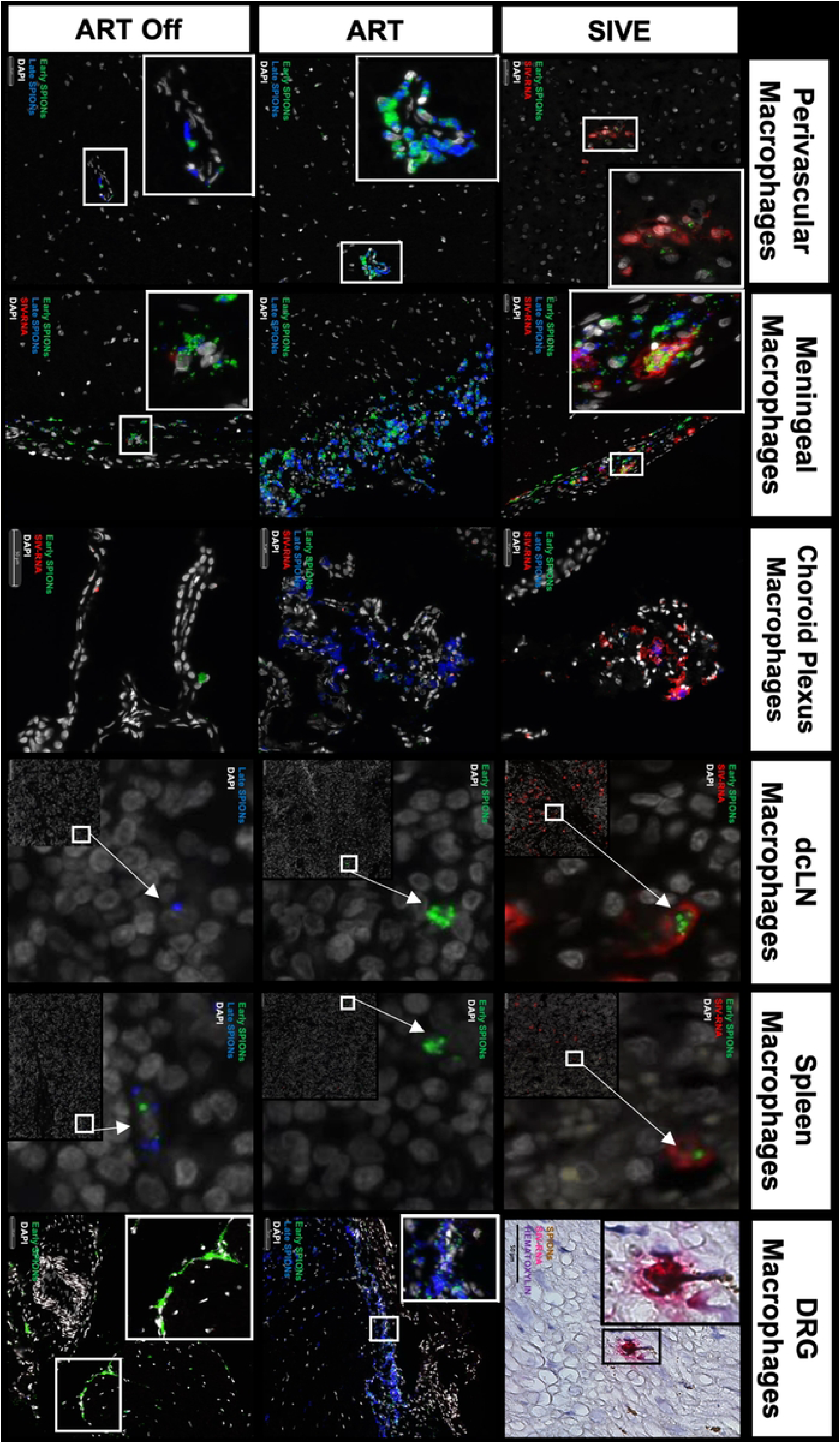
SPION+ SIV-RNA+ macrophages in the parenchyma, meninges, choroid plexus, dcLN, spleen, and DRG of animals with AIDS and SIVE. SIV-RNA (red) was detected in the CNS perivascular space, meninges, choroid plexus, and outside of the CNS in the dcLN, spleen, and DRG using RNAscope in situ hybridization. Early (green) SPION were i.c. injected 12-14 dpi and Late (artificially colored blue) SPION were i.c. injected 30 days prior to sacrifice. Images were captured from scanned sections. Nuclei are stained with Dapi (grey).

**Figure 4.**
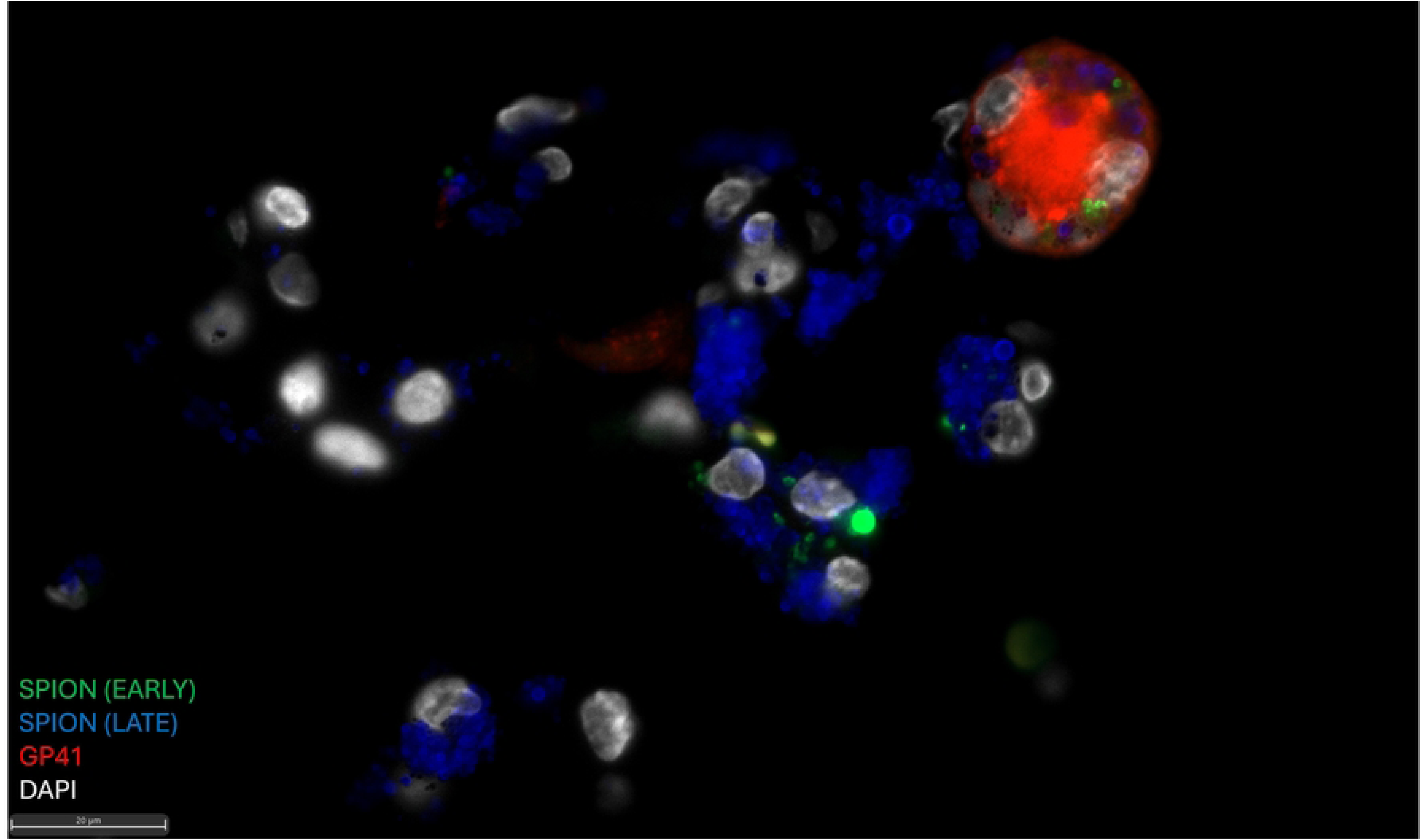
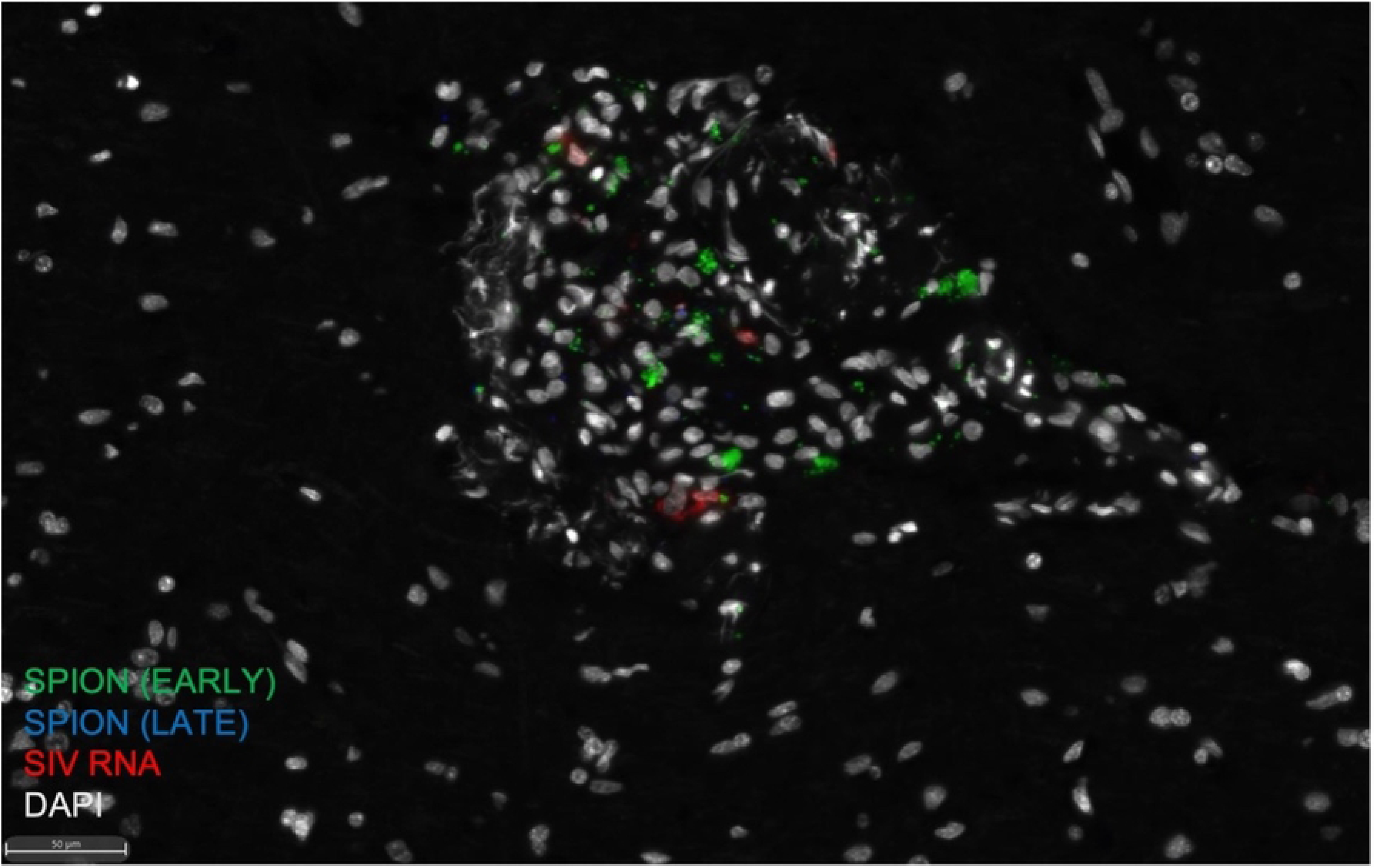
GP41+ SPION+ MNGC in the CNS of an animal with AIDS and SIVE. **(A)** SPION (Green and Blue) localizes to a gp41+ (red) MNGC in the cerebellum. **(B)** SPION (Green and Blue) localizes to a SIV-RNA+ (red) macrophages within lesions in the CNS cortical tissue. Nuclei are stained with Dapi (grey).

**Table 2.**
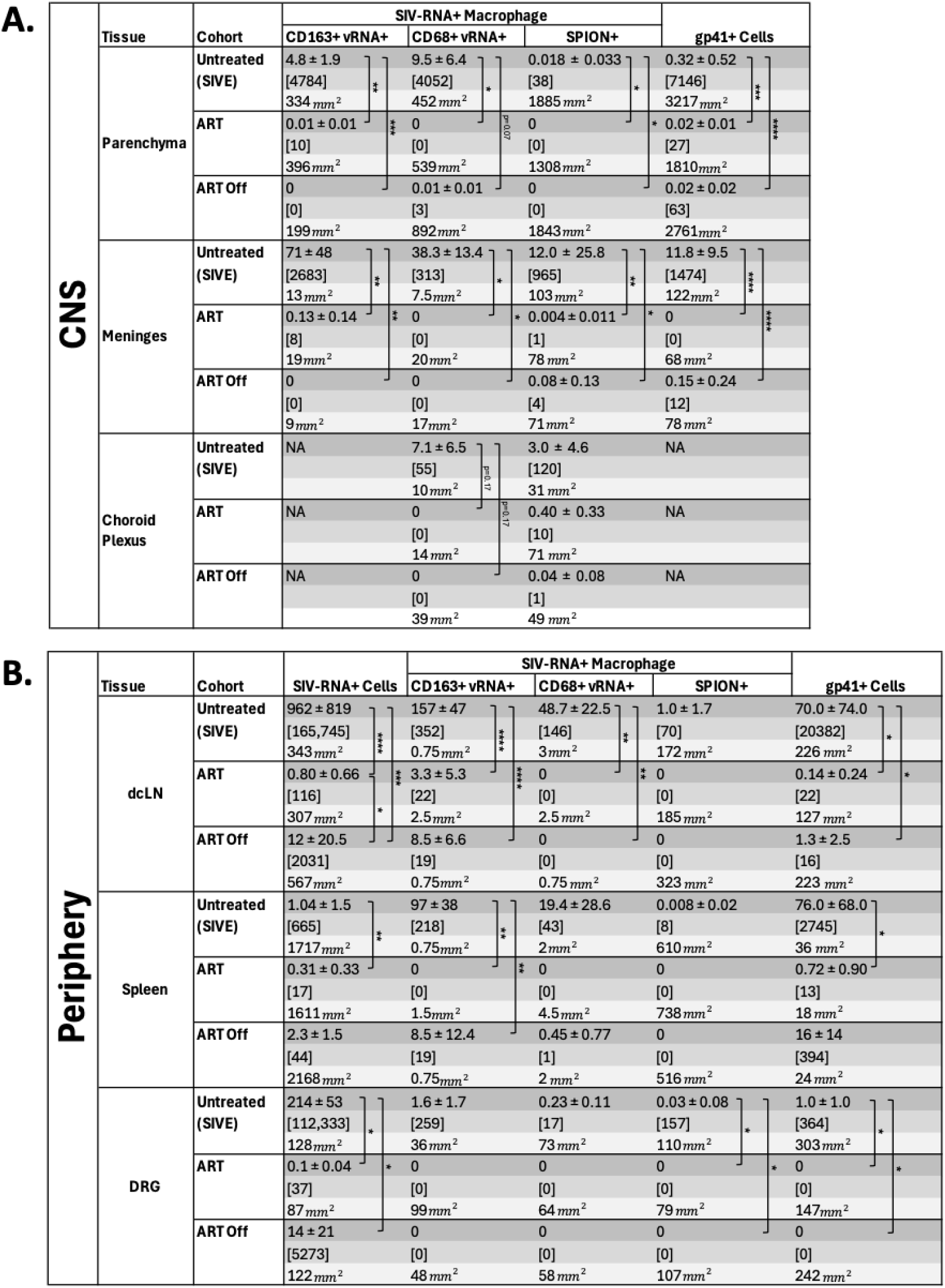
SPION+ SIV-RNA+ macrophages and gp41+ cells are reduced with ART but not in the meninges and choroid plexus which rebound following ART interruption.

SIV-RNA+ macrophages and cells were detected using ultrasensitive SIV-RNAscope with double-labeled CD163+ and CD68+ and GP41+ was detected using IHC. Whole tissue sections were analyzed and the number of positive cells assessed using fluorescence. A minimum of 2 CNS cortical regions and an average of 3 peripheral tissue sections were analyzed per animal. Counts are the mean ± SEM of the number of viral RNA+ cells, macrophages, SPION-labeled viral RNA+ macrophages, and gp41+ cells. Prism was used for graphing and statistical analysis Kruskal-Wallis nonparametric test was followed by Dunn’s post-hoc test, ∗P < 0.05, ∗∗P < 0.01, and ∗∗∗∗P < 0.0001.

Outside the CNS, there are significantly increased numbers of SPION+ SIV-RNA+ and SIV- RNA+ macrophages in AIDS and SIVE animals that are significantly reduced with ART (p<0.05) and do not rebound with ART interruption **(Table 2)**. There are few scattered CD163+ and CD68+ SIV-RNA+ macrophages in the dcLN, spleen, and DRG in SIVE animals that are eliminated with ART and did not rebound following ART interruption. **(Table 2)**. Overall, the data are consistent with ART reducing the number of productively infected and SPION+ virally infected macrophages in the CNS and periphery, and productive SIV-RNA+/gp41+ numbers rebound in the meninges, choroid plexus, dcLN, and spleen but not in CNS perivascular macrophages with ART interruption.

In parallel to the above studies, we assessed cell-associated SIV-RNA and DNA in FACsorted CD14+ monocytes and plasma virus longitudinally. Overall, with ART, plasma virus load decrease 4.5 logs to approximately 1x10^2^_log10_copies/ml (Figure 1b). Plasma viral load does not decrease to undetectable levels because of the short length of time animals are on ART and likely also because they are CD8 lymphocyte depleted. At 21 days p.i. there is an average of ∼6.9 x 10 ^4^ SIV-vRNA and ∼7.6 x 10^1^ SIV-vDNA copies per 10^6^ monocytes (n=8). At the same time point, with ART, cell-associated SIV-RNA and DNA in CD14+ monocytes decreased from ∼2.5 x 10^5^ SIV-vRNA and ∼1.9 x 10^3^ SIV-vDNA copies per 10^6^ monocytes to 4.0 x 10^4^ SIV-vRNA and 7.5 x 10^3^ SIV-vDNA copies per 10^6^ monocytes (n=4). At necropsy, non ART animals with AIDS and SIVE animals had ∼4.4 x 10^5^ SIV-vRNA and ∼9.7 x 10^3^ SIV-vDNA copies per 10^6^ monocytes, and undetectable monocyte SIV-RNA and -DNA in ART animals (n=4). Four weeks following ART interruption, at necropsy, monocyte associated SIV-RNA and DNA rebounded to ∼5.2 x 10^3^ SIV-vRNA and 3.7 x 10^1^ SIV-vDNA copies per 10^6^ monocytes (n=4).

### ART reduces early SPION+ perivascular macrophages in the CNS that corresponds to an increase in EARLY SPION+ macrophages in the periphery

To better define the timing and establishment of CNS macrophage viral reservoirs and the effect of ART and ART interruption on CNS macrophage retention and traffic out, we injected two different colored SPION. Green, fluorescent SPION were injected 12-14 days post-infection (Early), and red fluorescent SPION were injected 30 days prior to sacrifice (between 3-28 days prior to sacrifice, Late) **(Table 1**, **Figure 1)**. Our preliminary studies and work by others demonstrated SPION are internalized by macrophages within 30 minutes and remain intact in animals for greater than 120 days or the duration of our experiments (54, 55). We analyze the labeled cells in terms of ones that have early, late, or dual-labeled SPION. There are significantly (p<0.0001) greater numbers of early versus late and dual-SPION+ perivascular macrophages in AIDS and SIVE animals, ART, and ART interruption animals **(Figure 5A**. **Table 3)**. In contrast, there are equivalent numbers of early, late, and dual-labeled meningeal macrophages in SIVE, ART, and ART interruption animals **(Figure 5A&B. Table 3)**. The distribution of early and late SPION+ choroid plexus macrophages is similar to the meninges with equivalent numbers of early, late, and dual SPION+ macrophages. There are slightly greater numbers of early (SIVE: 3-fold, ART 4-fold, ART Off 4.6-fold) versus late or dual choroid plexus macrophages **(Table 3)**. Overall, these data suggest the CNS perivascular macrophage viral reservoir is established early in infection suggesting that meningeal and choroid plexus macrophages have ongoing recruitment of macrophages that increases with the development of AIDS and SIVE.

**Figure 5.**
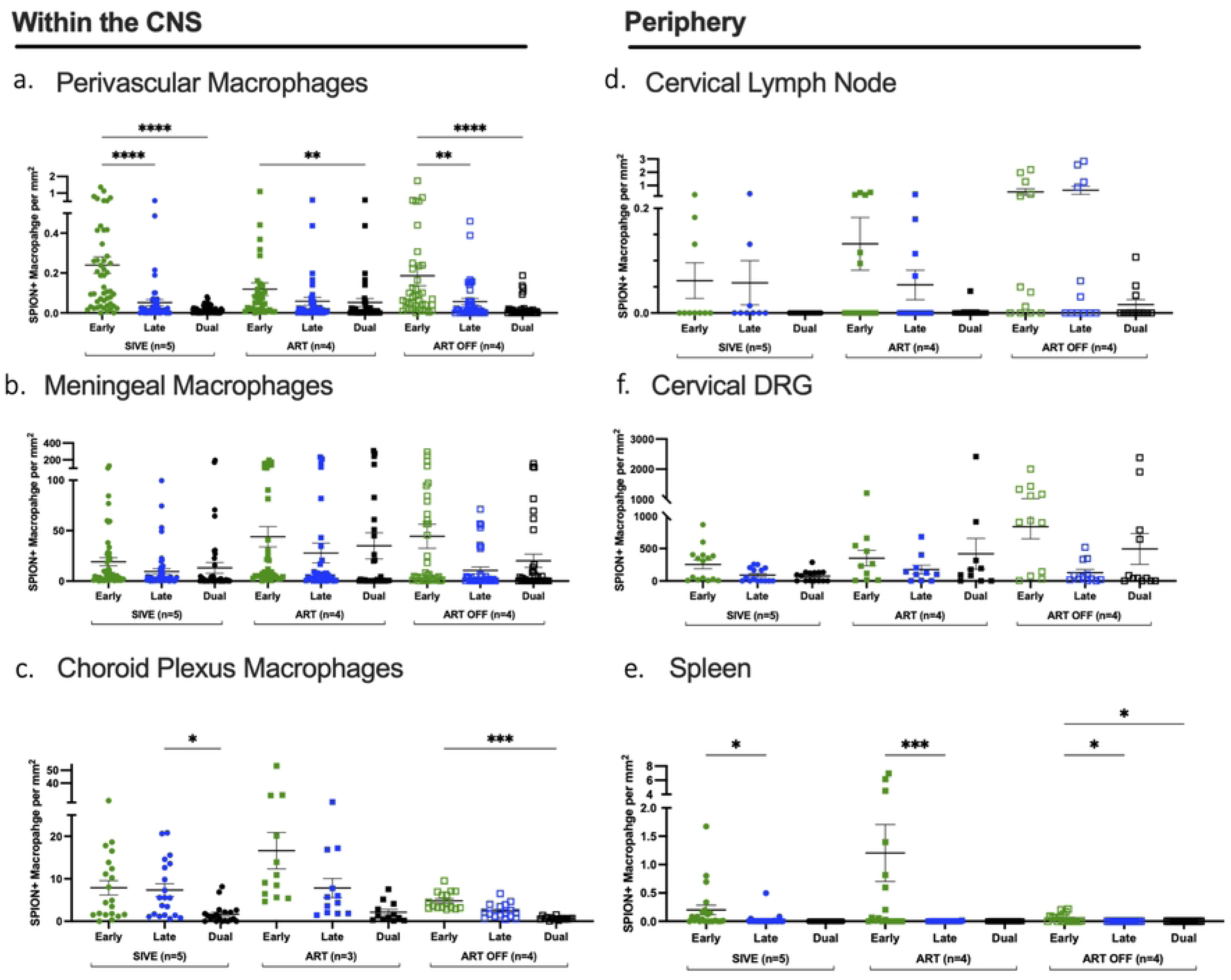
SPION-containing macrophages in and outside of the CNS are primarily labeled with SPION from early injection. Distribution of early, late, and dual-labeled SPION-labeled macrophage within the CNS **(A)** parenchyma, **(B)** meninges, **(C)** choroid plexus, and in the periphery including **(D)** dcLN, **(E)** spleen, and **(F)** DRG. Data points are the number of positive cells detected in one tissue section per animal and a minimum of 2 tissue sections were analyzed per animal. Prism was used for graphing and statistical analysis Kruskal-Wallis nonparametric test was followed by Dunn’s post-hoc test, ∗P < 0.05, ∗∗P < 0.01, and ∗∗∗∗P < 0.0001.

**Table 3.**
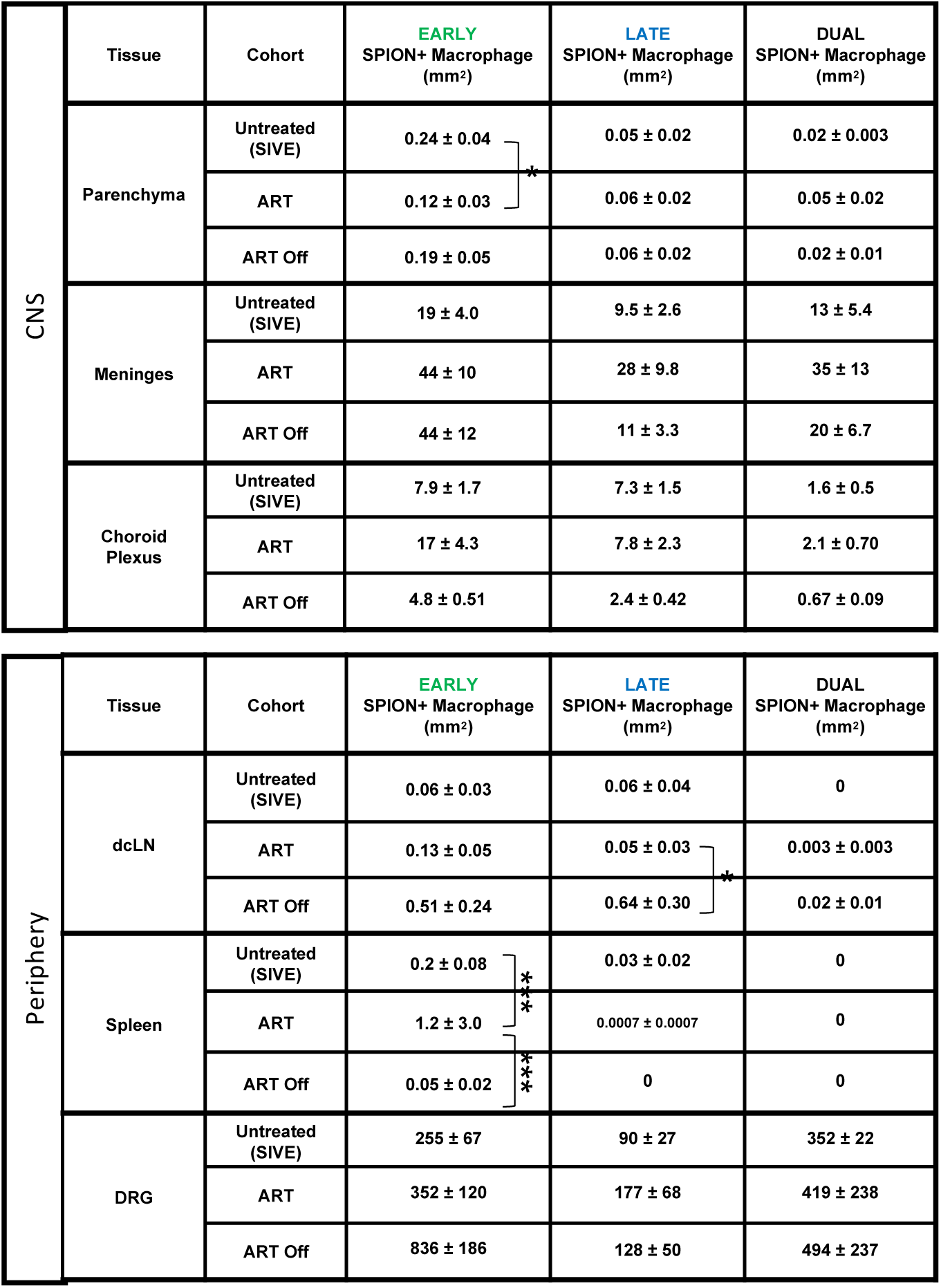
Distribution of early, late, and dual SPION+ macrophage in the CNS and periphery.

Whole tissue sections were analyzed, and the number of positive cells were identified using fluorescence. A minimum of 2 CNS cortical tissues were analyzed and an average of 3 peripheral sections were examined per animal, per tissue region. Counts are the mean ± SEM of the number of SPION+ macrophages (Early, Late, or Dual (both early and late) SPION). SIVE, SIV encephalitis; dcLN, deep cervical lymph node; DRG, dorsal root ganglia.

### ART animals have greater numbers of early SPION+ macrophages in the periphery compared to AIDS and SIVE animals

Outside of the CNS in the dcLN, spleen, and DRG, there is a trend of greater numbers of early than late or dual SPION+ macrophages with SIVE, ART, and ART interruption although these did not reach statistical significance **(Figure 5C**. **Table 3)**. With ART, there are greater numbers of early SPION+ macrophages in the spleen and few to no late or dual SPION+ macrophages **(Figure 5D**. **Table 3)**. In the DRG, there are greater numbers of early and dual SPION+ macrophages compared to late in animals with AIDS and SIVE, ART, and following ART interruption **(Figure 5E**. **Table 3)**. We next analyzed the number of early, late, and dual SPION+ virally infected macrophages in the CNS and periphery. The SPION+SIV-RNA+ perivascular macrophages in SIVE animals are primarily labeled with early SPION, that are eliminated with ART, and do not rebound with ART interruption **(Table 4)**. In contrast to perivascular macrophages, SIVE animals have greater numbers of Dual SPION+ SIV-RNA+ meningeal macrophages than early or late SPION **(Table 4)**. SPION+ SIV-RNA+ choroid plexus macrophages are primarily labeled with late SPION that are not eliminated with ART **(Table 4)**. The majority of SPION in SIVE animals with SIV-RNA+ macrophages in the dcLN are late inoculated **(Table 4)**. The SPION+SIV-RNA+ macrophages in the dcLN are eliminated with ART and do not rebound with ART interruption **(Table 4).** In the spleen of SIVE animals, there are equivalent numbers of early and late injected SPION+SIV-RNA+ macrophages that are eliminated with ART and do not rebound following ART interruption **(Table 4).** These data support the early establishment of the CNS macrophage viral reservoir, ART resolving CNS inflammation with subsequent traffic of macrophages out early, and increased traffic of virally infected macrophages out of the CNS that occurs late during infection with AIDS and SIVE.

**Table 4.** ART eliminates SPION+SIV-RNA CNS perivascular macrophages and SPION+SIV-RNA macrophages outside of the CNS but SPION+SIV-RNA+ meningeal and choroid plexus macrophages persist.

|  | Cohort | Tissue | SIV-RNA+SPION+ Macrophage (mm<sup>2</sup>) |  |  |
| --- | --- | --- | --- | --- | --- |
|  |  |  | EARLY | LATE | DUAL |
| CNS | Untreated (SIVE) | Parenchyma | 0.012 ± 0.027 [22] | 0.005 ± 0.008 [13] | 0.002 ± 0.003 [3] |
|  |  | Meninges | 2.1 ± 3.0 [191] | 3 ± 6 [250] | 7 ± 17 [524] |
|  |  | Choroid Plexus | 0.61 ± 0.62 [23] | 1.9 ± 3.5 [77] | 0.50 ± 0.68 [20] |
|  | ART | Parenchyma | 0 [0] | 0 [0] | 0 [0] |
|  |  | Meninges | 0 [0] | 0.004 ± 0.01 [1] | 0 [0] |
|  |  | Choroid Plexus | 0 [0] | 0.29 ± 0.52 [7] | 0.18 ± 0.27 [3] |
|  | ART OFF | Parenchyma | 0 [0] | 0 [0] | 0 [0] |
|  |  | Meninges | 0.01 ± 0.04 [1] | 0.01 ± 0.04 [1] | 0.03 ± 0.07 [2] |
|  |  | Choroid Plexus | 0 [0] | 0 [0] | 0.042 ± 0.084 [1] |
| Periphery | Cohort | Tissue | SIV-RNA+SPION+ Macrophage (mm<sup>2</sup>) |  |  |
|  |  |  | EARLY | LATE | DUAL |
|  | Untreated (SIVE) | dcLN | 0.07 ± 0.14 [20] | 0.9 ± 1.7 [50] | 0 [0] |
|  |  | Spleen | 0.0025 ± 0 [3] | 0.004 ± 0.009 [5] | 0 [0] |
|  |  | DRG | NA | NA | * 0.03 ± 0.086 [157] |
|  | ART | dcLN | 0 [0] | 0 [0] | 0 [0] |
|  |  | Spleen | 0 [0] | 0 [0] | 0 [0] |
|  |  | DRG | NA | NA | 0 [0] |
|  | ART OFF | dcLN | 0 [0] | 0 [0] | 0 [0] |
|  |  | Spleen | 0 [0] | 0 [0] | 0 [0] |
|  |  | DRG | NA | NA | * 0.0012 ± 0.008 [3] |

SIVE, SIV encephalitis; dcLN, deep cervical lymph node; DRG, dorsal root ganglia. Counts are the mean ± SEM of the number of viral RNA+ cells and SPION-labeled viral RNA+ macrophages.

Whole tissue sections were analyzed to obtain the number of positive cells based on the detection of fluorescence. A minimum of 2 cortical CNS tissues and an average of 3 peripheral sections were examined per animal, per tissue region.

## Discussion

Few studies have investigated CNS macrophage accumulation and traffic out normally, with HIV and SIV infection, or ART, and ART interruption. Using i.c. injection of SPION, that labeled CNS perivascular, meningeal, and choroid plexus macrophages we found animals with AIDS and SIVE had increased CNS SPION+ macrophages consistent with CNS macrophage accumulation. There was a differential distribution of SPION+ macrophages with 50 to 200-fold greater number of SPION+ meningeal macrophages than SPION+ perivascular macrophages, and 1 to 5-fold greater number of meningeal than choroid plexus macrophages. These results are consistent with greater numbers of total meningeal versus perivascular macrophages and interestingly, the ratio of total macrophages versus SPION+ macrophages in the two compartments did not change between SIVE, ART, and ART Off animals. ART reduced the number of SPION+ perivascular macrophages within the CNS where AIDS and SIVE animals have approximately 4x more SPION+ perivascular macrophages than ART animals. ART did not affect the number of SPION+ macrophages in the meninges and choroid plexus. Following ART interruption, there were increased numbers of SPION+ meningeal macrophages and a decrease of SPION+ choroid plexus macrophages compared to ART animals. Increased numbers of SPION+ macrophages in the meninges with ART interruption may account for decreased choroid plexus macrophages where increased traffic of BM macrophages from the choroid plexus to the CSF could result in a corresponding increase in the SPION+ meningeal macrophages. We note that SPION+ macrophages in the perivascular space, dcLN, and the spleen typically contain fewer SPION per cell than macrophages in the meninges and DRG (data not shown). The differential numbers are likely due to the fluid-SPION interface in the different macrophage compartments (18, 56, 57). Additionally, the rate of turnover of these cells normally, and once they have taken up SPION is also likely a factor. It is possible that once a macrophage is sufficiently loaded with SPION, certain egress routes, such as passing through the cribriform plate, are less feasible. SPION uptake by macrophages could ultimately slow their exit from the CNS and potentially contribute to the accumulation of these cells in different compartments as we observed in the meninges. Alternatively, macrophages that are more able to traffic out of the CNS, might do so once taking up fewer SPION. An assumption of our studies, supported by the work of others, is that SPIONs unlike dextrans (33, 47, 58–60) do not leak out of the CNS. This is supported by the size of SPION used in our studies and also our lack of finding single, non-cell associated SPION inside and outside the CNS. It is a valid question whether SPION are broken down and exit the CNS as others have previously shown (61, 62), though this is unlikely in our timeframe examined (<1 year) and beads used due to the stability of their encapsulated synthetic polymer coating (60, 63–65). A more likely outcome in our study is the clearance of SPIONs through the kidneys or the induction of iron-programmed cell death due to high levels of iron ions and ROS, leading to lipid peroxidation and eventual cell death (60, 63–66). Overall, these data are consistent with a differential turnover of CNS perivascular, meningeal, and choroid plexus macrophages and a reduction primarily in CNS perivascular macrophages with ART. Lastly, that macrophages exit particular tissues and traffic to the dcLN has recently been demonstrated in rodent models (47, 48, 67) and likely also occurs in the CNS, but hasn’t been studied to date.

Macrophage turnover in the CNS has been shown to occur normally at a rate that increases with inflammation(38). Less well known is the effect of ART and the contribution of macrophage accumulation with inflammation early and late with regard to viral infection. Using two different colored SPION injected early and late, we found greater numbers of early versus late or dual labeled macrophage in all tissues examined regardless of treatment. Perivascular macrophages are primarily labeled with early SPION suggesting they are a resident, low-turnover macrophage population with SIV infection. A greater number of early SPION-labeled macrophages may occur due to a limited capacity for SPION uptake. Regardless, this data is consistent with our previous reports of early recruitment of CD163+ BM-derived perivascular macrophages to the CNS with SIV infection (33, 68, 69) and we observed that perivascular macrophage turnover occurs at a low rate that is increased with SIV infection (12, 38, 41, 68, 69). In contrast, meningeal macrophages had equivalent numbers of early and late injected SPION+ macrophages in SIVE, ART, and following ART interruption, suggesting these are a more static locally maintained macrophage population that is unchanged with ART or following ART interruption. Fluid dynamics in regions such as the meninges and even DRG may contribute to the even distribution of early, late, and dual-labeled macrophages in these compartments, as these cells are continuously exposed to CSF-containing SPION. Choroid plexus macrophages also had equivalent numbers of Early and Late but few dual SPION+ macrophages in SIVE, ART, and ART interruption animals, consistent with a high rate of traffic and turnover of monocytes/macrophages. The continuous turnover and exit of CNS macrophages to the periphery may occur with CNS lesion resolution and provide a mechanism to prevent chronic inflammation and tissue damage with SIV and HIV infection, similar to findings in recent rodent studies(70, 71). Overall, our data using SPION to label CNS macrophages to investigate turnover shows the traffic of CNS macrophages out to the periphery (dcLN, spleen, and DRG). In the periphery, we typically observed more early-labeled macrophages, which may indicate a delay in the clearance of SPION+ cells from the CNS rather than the rapid drainage of free SPIONs. Additionally, this increase in the number of early labeled SPION+ macrophages in the periphery could be consistent with the traffic of a DC-like macrophage that exit the CNS possibly from the perivascular space, similar to what others have shown in rodents(47, 48). In this study we did not directly measure traffic of specific subpopulation of SPION+ CNS macrophage out of the CNS nor did we define specific pathways of exit. Furthermore, specific studies would be required to better understand where macrophage populations leave the CNS. Such future studies are likely critical to understanding CNS inflammation and the resolution of inflammation. This may be especially critical in human diseases like multiple sclerosis and HIV infection with ART.

Pathways by which macrophages can exit the CNS have been proposed and described(19, 23, 46, 47, 51, 70–84), including i) dorsal and basal dural lymphatics ii) the olfactory bulb-cribriform lymphatic axis, ii) the choroid plexus, and ii) egress via spinal pathways (50, 76, 85). More recently, discontinuities in the arachnoid barrier at bridging veins are another path of immune cell and CSF drainage into the dura and egress via dural lymphatics (86). Perivascular and meningeal macrophages could exit with CSF via meningeal lymphatics or cranial nerve routes to reach the dcLN (49, 50, 87). We find increased numbers of SPION+ macrophages in the dcLN which likely exited the CNS via nasal and meningeal routes rather than in the superficial cLN which would likely be drainage for cranial nerve egress (49, 88). As the choroid plexus macrophages interface blood and CSF on the apical side, stromal macrophages can exit via perivenous routes and/or migrate through the choroid plexus to the CSF then through the subarachnoid space. Choroid plexus Kolmer macrophages likewise could leave via pathways of CSF drainage (19, 47, 48, 75). Additionally, immune cells could exit the CNS via CSF drainage pathways in the spine, including arachnoid villi to spinal veins (89–91), along spinal nerve roots (DRG) to epidural lymphatics (92, 93), and routes through the arachnoid layer to the spinal meninges to dural lymphatics (94, 95). The egress of macrophages, some of which could be infected, is crucial for understanding CNS reservoir resolution and possible redistribution of CNS virus.

An important consideration is whether CNS SIV or HIV virally infected macrophages can traffic out of the CNS to the periphery. We found SIVE animals with SIV-RNA+ SPION+ perivascular macrophages, meningeal, choroid plexus macrophages, and SIV-RNA+ SPION+ macrophages in the dcLN, spleen, and DRG. With ART, SIV-RNA+ SPION+ macrophages were eliminated in the CNS parenchyma (perivascular macrophages) and periphery, but not meninges and choroid plexus where we found scattered SIV-RNA and gp41+ macrophages that persisted with ART. With ART interruption, there was a rebound of SIV-RNA+ SPION+ and SIV- RNA+ meningeal macrophages, choroid plexus macrophages, macrophages in the spleen, and plasma virus but importantly not SIV-RNA+SPION+ perivascular macrophages. We did not find any SIV-DNA+ macrophage with SPION within or outside the CNS with ART which is most likely due to the reduced sensitivity of the DNAscope assay compared to SIV-RNAscope assay in our hands, although we note in the monocyte associated SIV- RNA and -DNA, we consistently find lower levels of DNA. Clements et al. recently determined that virologically suppressed people living with HIV have persistent, replication-competent virus in monocyte-derived macrophages in blood using a quantitative viral outgrowth assay (11). Although we did not find SIV- RNA+SPION+ macrophages with ART in the peripheral tissue sections, Clements and colleagues also showed that despite low levels of viral DNA, replication-competent virus persists in monocyte-derived macrophages despite ART(11). Importantly, with ART interruption, we find a rebound of virally infected macrophages in CNS meninges, choroid plexus, plasma virus, and spleen, but not in the CNS perivascular macrophages or cell-associated SIV-RNA and DNA in PBMCs. These data suggest that with AIDS and SIVE there can potentially be ongoing reseeding of the periphery from the CNS viral reservoir that is eliminated with ART and with ART interruption virus rebounds from the plasma, meninges, and spleen. As macrophages in the choroid plexus have direct access to egress to blood, there may be a contribution to viral rebound from this CNS compartment with ART interruption as we found to occur in our study. While the CNS has been historically regarded as an immune-privileged site with restricted immune cell trafficking and controlled egress of immune cells—particularly macrophages—is critical for controlling inflammation. Our data underscores the dynamic nature of macrophages in the CNS that can migrate from the BM to the CNS and traffic antigens out of the CNS with inflammation or viral infection. The egress of macrophages from the CNS with ART occurs with and likely results in the resolution of inflammation and reduced macrophage infection. While macrophage traffic out of the CNS challenges the conventional notion of CNS immune privilege, it is critical for the regulation of neuroinflammation.

Using two different colored, fluorescently labeled SPION i.c. injected early (12-14 days post-infection) and late (30 days prior to sacrifice) with SIV-RNAscope we studied the timing and location of CNS macrophage viral reservoir establishment. We found significantly more early SPION+ SIV-RNA+ perivascular macrophages compared to late or dual in AIDS and SIVE animals that were eliminated with ART and did not rebound with ART interruption. This suggests and agrees with our previous data(33) that the CNS perivascular macrophage viral reservoir is established during early infection. Conversely in the meninges, we found equal distribution of SIV-RNA+ early, late, and dual-labeled SPION+ macrophages in animals with AIDS and SIVE that were eliminated with ART but rebounded with ART interruption. This suggests ongoing recruitment of SIV-RNA+ macrophages in the meninges with AIDS and SIVE that is eliminated with ART and rebounds following ART interruption. In the choroid plexus, we found greater numbers of Late SPION+ SIV-RNA+ macrophages in SIVE animals that were not eliminated with ART. This data is consistent with the choroid plexus being a site of entry for BM-derived monocytes to the CNS with a high rate of turnover that is increased with SIV infection (33, 38). It is important to note in CD8-depleted macaques, we do not reduce plasma viral load to undetectable levels with ART (1x10^4^ log_10_ copies/mL). Thus, it would be expected to find viral RNA+ monocytes in the blood, possibly even containing SPION (not examined). These data support the early establishment of the CNS macrophage viral reservoir, ART resolving CNS inflammation by inducing traffic out early, differential rates of viral seeding in the parenchyma, meninges, and choroid plexus, and increased traffic of virally infected SPION+ macrophages out to the periphery with AIDS.

One site of particular interest to the CNS with regards to neuroimmune modulation is the choroid plexus as it is viewed as an immune cell gateway to the brain (22, 26, 27, 36, 96). HIV and SIV-infected cells are consistently demonstrated in the choroid plexus(26, 27, 97) but whether the choroid plexus is a viral reservoir has not been established. We found an accumulation of SPION+ macrophages within the choroid plexus, which could have been labeled via diffusion of SPION with CSF across the epiplexus, recirculation of SPION or SPION+ monocytes/macrophage. Previously, using an anti-A4B1 antibody (Tysabri) that blocks the traffic of monocytes and macrophages to the CNS, we prevented the establishment of the CNS viral reservoir and reduced neuronal injury (43). Because A4 integrin is expressed on both sides of the choroid plexus, anti-A4B1 treatment could prevent cellular traffic to both the blood and CSF from the choroid plexus and determine if there is recirculation of monocytes/macrophages to the CNS. By blocking the traffic of immune cells out of the CNS by Visudyne ablation of meningeal lymphatics, Louveau *et al.* found there is a reduction in clinal symptoms from EAE while conversely improving the function of meningeal lymphatics by increasing the diameter with vascular endothelial growth factor C (VEGF-C) and subsequently increasing traffic of immune cells out of the CNS in aged mice improved clinical symptoms (19, 46–51). Our SPION data indicates that traffic of CNS macrophages out to the dcLN, spleen, and DRG occurs normally with SIV infection and is increased with ART concurrently with the resolution of CNS inflammation. Using anti-A4B1 antibody in conjunction with ART to block the accumulation of macrophages in the CNS could prevent viral recrudescence of SPION+ macrophages we found in the CNS meninges and choroid plexus with ART interruption.

The results of this study suggest myeloid traffic within and outside the CNS with HIV and SIV infection plays a major role in the resolution of CNS inflammation. Our study found that animals with SIVE had more SPION+ macrophages in the CNS compared to those without infection or those treated with ART. In contrast, the ART animals had fewer SPION+ CNS macrophages but more SPION+ macrophages that moved out of the CNS to the periphery, concurrent with the resolution of CNS inflammation. The traffic of CNS viral reservoir out of the CNS suggests a paradox because historically, the CNS has been seen as a place where the immune system is restricted, and the movement of immune cells, especially macrophages, is tightly controlled to manage inflammation. Moreover, it is generally considered that ART clears HIV and SIV in the CNS but it is possible that virally infected macrophages in the CNS can persist and traffic out to the periphery to redistribute virus. Our findings emphasize that macrophage recruitment to the CNS is dynamic as they migrate from the bone marrow (BM) and respond to CNS inflammation and viral infections and may eventually traffic out.

## Methods

### Ethics Statement

All animal work was approved by the Tulane National Primate Research Center Care and Animal Use Committee. The TNRPC protocol number is 3497 and the animal welfare assurance number is A4499-01.

### Animals and Viral Infection

A total of 20 adult rhesus macaques (*Macaca mulatta*) born and housed at the Tulane National Primate Research Center in strict adherence to the “Guide for the Care and Use of Laboratory Animals” were used in this study **(Figure 1A**. **Table 1)**. CD8+ lymphocytes were depleted to achieve rapid AIDS (3 to 4 months) with >75% incidence of SIVE, as previously described (33, 34, 68, 98). CD8+ T lymphocyte depletion was monitored longitudinally by flow cytometry and all macaques were persistently depleted (>28 days). Animals were experimentally infected intravenously (i.v) by inoculation with a bolus of SIVmac251 viral swarm (20 ng of SIV p28) provided by Dr. Ronald Desrosiers, over 5 minutes. At 21 days post-infection, n=14 macaques began a 12 - 15-week regimen of antiretroviral therapy (ART) consisting of Raltegravir (Merck, 22 mg/kg) given orally twice daily, and Tenofovir (Gilead, 30 mg/kg) and Emtricitabine (Gilead, 10 mg/kg) combined in a sterile solution given once-daily, s.c. Ten animals were euthanized on ART and 4 were removed from ART for 4 weeks to allow for viral rebound. Animals were euthanized based on the recommendations of the American Veterinary Medical Association Guidelines for the Euthanasia of Animals upon developing signs of AIDS, which included: a >15% decrease in body weight in 2 weeks or >30% decrease in body weight in 2 months; documented opportunistic infection; persistent anorexia>3 days without explicable cause; severe, intractable diarrhea; progressive neurological symptoms; or significant cardiac or pulmonary symptoms. SIV encephalitis (SIVE) was defined by the presence of multinucleated giant cells (MNGC) and the accumulation of macrophages (68). Longitudinal plasma viral load (PVL) was assessed as previously described (99–101) to monitor viral suppression during treatment and rebound following ART interruption **(Figure 1B)**. For PVL, 500*μL* of EDTA plasma was collected and plasma virions were pelleted by centrifugation (20,000 x g for 1 hour). The sensitivity threshold of the assay was 100 copy Eq/mL with an average intra-assay coefficient variation of less than 25%. Log-transformed PVLs below the limit of detection were set to 0 for statistical analysis.

### SPION (and Dextrans)

Superparamagnetic iron oxide nanoparticles (SPION) were obtained from Bang Laboratories Inc. (PS-COOH Mag/Encapsulated, MEDG001, and MEFR001, Fishers, IN). SPION have internal fluorescence (Dragon Green [480/520] and Flash Red [660/690]) and are iron oxide nanospheres encapsulated in an inert polymer with an average particle size of 0.86*μm* diameter. SPION were prepared in a class 2 biosafety cabinet. 3 mL of stock solution and 7 mL of sterile, low endotoxin 1XPBS were added to a 15 mL conical tube and gently mixed. A magnet was used to separate the SPION from the liquid and an additional 7 mL of 1XPBS was added for a second wash. This process was repeated 7 - 10 times to ensure all original buffer was removed and the SPION were resuspended in a sterile 1XPBS solution. SPION reconstituted in 1mL of sterile 1XPBS, at a final concentration of 33 mg/mL. In animals receiving fluorescent Dextran (RITC-Dextran,10 KD, Invitrogen, CA) was administered at a concentration of 33 mg/mL.

### Intracisternal Inoculation Procedure

To avoid an increase in intracranial pressure, 1 mL of cerebrospinal fluid (CSF) was removed from the cisterna magna prior to the inoculation of SPION or RITC-Dextran. Dragon Green [480/520] SPION (33 mg/mL) were administered 2 weeks post-inoculation at 12 or 14 dpi. Flash Red [660/690] SPION (33 mg/mL) were injected late during infection prior to euthanasia (Range 3-22 days).

### Tissue Collection and Processing

Animals were anesthetized with ketamine-HCl and euthanized with i.v. pentobarbital overdose and exsanguinated. Blood was collected and Heparin Sulfate was administered i.v. and given 5 minutes to diffuse. Sodium pentobarbitol was administered via intracardiac stick and CSF was collected. Following CSF collection, animals underwent perfusion with 3L of chilled 1XPBS. Postmortem examination was performed by a veterinary pathologist that confirmed the presence of AIDS-defining lesions as previously described (33, 34, 68, 98). Brain (frontal, temporal, occipital, parietal, and choroid plexus) and peripheral tissues (cervical lymph nodes, spleen, and DRG) were: i) collected in zinc-buffered formalin and embedded in paraffin, ii) fixed with 2% paraformaldehyde for 4-48 hours, sucrose protected and embedded in optimal cutting temperature (OCT) compound for SPION analysis, or iii) snap frozen in OCT without fixation (spleen).

### Immunohistochemistry and Immunofluorescence

IHC was performed as previously described using the antibodies targeting CD163 (1:250, Leica (Deer Park, IL)), CD68 (1:100, DAKO (Carpinteria, CA)), CD206 (1:1000, R&D Systems (Minneapolis, MN)), and IBA1+ macrophages (1:100, Wako (Osaka, Japan)) (33, 34, 98). Briefly, formalin-fixed paraffin-embedded sections were deparaffinized and rehydrated followed by antigen retrieval with a citrate-based Antigen Unmasking Solution (Vector Laboratories, Burlingame, CA) in a microwave (900 W) for 20 min. After cooling for 20 min, sections were washed with Tris-buffered saline (TBS) containing 0.05% Tween-20 for 5 minutes before incubation with peroxidase block (DAKO, Carpinteria, CA) followed by protein block (DAKO, Carpinteria, CA) for 30 minutes and incubation with primary antibody. Following incubation with a peroxidase-conjugated polymer, slides were developed using a diaminobenzidine chromogen (DAKO, Carpinteria, CA) with Harris Hematoxylin (StatLab, McKinney, TX).

Immunofluorescence for CD163+, CD68+, IBA1+, and gp41+ macrophages was performed using antibodies and fluorochromes using CD163 (NCL-CD163 CE, AF568, Invitrogen (Carlsbad, CA)), CD68 (DAKO, AF568 (Carpinteria, CA)), IBA1+ macrophages (Wako, DISCOVERY OmniMap Anti-Rb HRP (Osaka, Japan)), and gp41+ (KK41+, 1:100, NIH (Manassas, VA)) on 2% paraformaldehyde (PFA) fixed frozen sections. 2% PFA, fixed frozen sections were thawed for 20 minutes at room temperature, unwrapped, submerged in a citrate-based Antigen Unmasking Solution, and microwaved for one minute and forty-five seconds and cooled to room temperature. Slides were permeabilized in a solution of phosphate-buffered saline with 0.01% Triton X-100 and 0.02% fish skin gelatin (PBS-FSG-TX100) followed by a PBS-FSG wash, transferred to a humidified chamber and blocked with 10% normal goat serum (NGS) diluted in PBS-FSG for 40 minutes, followed by a 60-minute primary antibody incubation, washes, and 40- minute secondary antibody incubation. Routine washes were performed and DAPI nuclear stain added for 10 minutes. Slides were mounted using a custom-formulated anti-quenching mounting media containing Mowiol (#475904, Calbiochem; San Diego, CA) and DABCO (#D2522, Sigma: St. Louis, MO) and allowed to dry overnight before being digitally imaged with a Zeiss Axio Scan.Z1. HALO software (HALO v3.4, Indica Labs; Albuquerque, NM) was used for quantification and analysis.

### Tissue Viral RNA and DNA Detection

Ultrasensitive SIV-RNAscope with probes for SIVmac251 was used to detect SPION-containing SIV-RNA positive cells within and outside of the CNS as previously described (98). Tissue sections were placed in a target antigen retrieval solution, heated, and treated with protease plus, and a hydrogen peroxide blocker according to the manufacturer’s protocol (Advanced Cell Diagnostics, Newark, CA). SIVmac239 RNAscope probes (Advanced Cell Diagnostics, Newark, CA) were hybridized at 40°C in the HybEZ II Hybridization System. The RNAscope 2.5 HD Assay amplification steps were applied according to the manufacturer’s protocol. Target RNA was visualized through the addition of chromogenic Fast Red A and Fast Red B (Advanced Cell Diagnostics, Newark, CA), and sections were counterstained with hematoxylin (Sigma-Aldrich) and mounted using Vectamount (Vector Laboratories). Viral RNA processed sections were subsequently stained for CD163+ or CD68+ macrophages using primary antibodies and IHC methods described above. Prussian blue iron staining and/or the internal fluorescence of SPION was used to detect vRNA+SPION-containing CD163+ macrophage. DNAscope was performed, as previously described (98, 102, 103). SIV-DNA was detected in situ using an SIV-DNA sense probe (Advanced Cell Diagnostics, Newark, CA) for RNAscope® Assay on 3 – 4 CNS cortical sections and 1 peripheral tissue (dcLN, spleen, and DRG) per animal. To reduce non-specific signal, brain tissues were pre-treated with 2N HCL for 30 min at room temperature.

### Detection and quantification of SPION-containing macrophages in tissues

SPION were detected in the central nervous system (CNS) and peripheral tissues by 1) light microscopy by morphology of the amber SPION beads, 2) Prussian blue iron staining (Sigma Aldrich Iron Stain, St. Louis, MO), and 3) internal fluorescence of Dragon Green or Flash Red. The number of SPION-containing macrophages with IHC for macrophages with CD163+ or CD68+ was counted. SPION+ macrophages in whole tissue sections were counted manually at 20x by light microscopy (Plan-Apochromat 620/0.7, Olympus; Japan) in a blinded fashion. Whole section tiling and stitching was done using a Zeiss Axio Imager M1 microscope (Zeiss; Oberkochen, Germany) with AxioVision (Version 4.8, Zeiss; Oberkochen, Germany) using Plan-Apochromat 620/0.8 and 640/0.95 Korr objectives followed by manual annotation of the parenchyma from the meninges into separate regions of interest (ROI) and tissue area reported as mm^2^.

For analysis of early versus late SPION+ macrophages and virally infected SPION+ macrophages, whole slide fluorescent images of stained sections were scanned using a Zeiss AxioScan Z.1 (Carl Zeiss MicroImaging, Inc., Thornwood, NY). Scanned sections were analyzed with HALO modular analysis software (HALO v3.4, Indica Labs; Albuquerque, NM). The parenchyma and meninges were first annotated into separate ROI/annotation layers and the number of early, late, and dual SPION-containing macrophages and vRNA-FISH early, late, and dual SPION-containing macrophages were counted using the FISH v.3.2.3 module (HALO v3.4, Indica Labs; Albuquerque, NM) and reported as the number of cells per mm^2^.

### Cell Associated Viral Load

A minimum of 100,000 T cells and Monocytes were isolated from frozen PBMCs using flow cytometry using a Sony SH800. Thawed PBMCs were incubated with fluorochrome-conjugated antibodies to isolate T cells and Monocytes including anti-C14-FITC (clone: HCD14, BioLegend), anti-CD3-PE (clone: UCHT1, BioLegend), anti-HLA-DR-ECD (clone: Immu-357, Beckman), anti-7- AAD-PECY5 (BD), anti-CD20-PERCPCY5.5 (clone: 2H7, BD Pharmingen), and anti-CD3-PECY7 (clone: SK7, BioLegend). Monocytes were selected based on size and granularity (FSC vs SSC) followed by selection of live (7-AAD^+^) CD14^+^HLA-DR^+^CD3^-^ CD20^-^ cells and T cells were likewise selected based on size and granularity (FSC vs SSC) followed by selection of live (7-AAD^+^) CD3^+^ HLA-DR^-^CD14^-^ cells. After sorting, cells were washed with 2% FBS-PBS and dry pelleted before cell-associated SIV viral load analysis was performed by The Quantitative Molecular Diagnostics Core at the Fredericks National Laboratory. Briefly, the Quantitative Molecular Diagnostics Core performed total DNA and RNA extraction from the sorted Monocytes and T cells and tested them separately using a hybrid assay format combining quantitative real-time standard curve interpolation and Poisson-based methods. Extracted RNA was resuspended and divided among 10 replicate quantitative polymerase chain reaction (qPCR) or quantitative reverse transcription polymerase chain reaction (qRT-PCR) reactions. SIV DNA testing was multiplexed with an assay for a single-copy genomic sequence from rhesus macaque CCR5 to normalize to cell numbers based on diploid genome equivalents. For specimens where all wells yielded positive PCR reactions, the SIV DNA copy number was calculated using the average of the standard curve interpolated values and normalized to cell numbers. When not all wells were positive, Poisson methods were used to calculate viral copy numbers, followed by normalization. SIV RNA values were determined by qRT-PCR and normalized to cell numbers based on CCR5 DNA copy numbers, accounting for near-quantitative RNA and DNA recovery with the employed extraction methods.

### Statistical Analysis

Statistical analyses were performed using Prism version 10.0 (GraphPad Software; San Diego, CA). Comparisons between animals with SIVE, ART, and following ART interruption were made using a nonparametric one-way analysis of variance (Kruskal-Wallis, GraphPad Software; San Diego, CA) with Dunn’s multiple comparisons. Statistical significance was accepted at p< 0.05 and all graphing was done using Prism (GraphPad Software; San Diego, CA).

## Acknowledgements

We would like to acknowledge the Core staff at the Tulane National Primate Research Center (RRID:SCR_008167) that worked on this project including the Anatomic Pathology Core (RRID: SCR_024606) Clinical Pathology Core (RRID: SCR_024609) Confocal Microscopy and Molecular Pathology Core (RRID: SCR_024613) (RRID:SCR_008167), and Virus Characterization, Isolation, Production and Sequencing Core (RRID:SCR_024679). We acknowledge Dr. Jeff Lifson, AIDS and Cancer Virus Program for monocytes associated SIV-RNA and DNA analysis.

## Abbreviations

ART: Antiretroviral therapy
CNS: Central Nervous System
HIV: Human Immunodeficiency Virus
SIV: Simian Immunodeficiency Virus
SPION: Superparamagnetic iron oxide nanoparticles
DRG: Dorsal Root Ganglia
AIDS: Acquired Immunodeficiency Syndrome
dcLN: Deep Cervical Lymph Node
SIVE: SIV-induced Encephalitis

